# Dopamine transients delivered in learning contexts do not act as model-free prediction errors

**DOI:** 10.1101/574541

**Authors:** Melissa J. Sharpe, Hannah M. Batchelor, Lauren E. Mueller, Chun Yun Chang, Etienne J.P. Maes, Yael Niv, Geoffrey Schoenbaum

## Abstract

Dopamine neurons fire transiently in response to unexpected rewards. These neural correlates are proposed to signal the reward prediction error described in model-free reinforcement learning algorithms. This error term represents the unpredicted or ‘excess’ value of the rewarding event. In model-free reinforcement learning, this value is then stored as part of the learned value of any antecedent cues, contexts or events, making them intrinsically valuable, independent of the specific rewarding event that caused the prediction error. In support of equivalence between dopamine transients and this model-free error term, proponents cite causal optogenetic studies showing that artificially induced dopamine transients cause lasting changes in behavior. Yet none of these studies directly demonstrate the presence of cached value under conditions appropriate for associative learning. To address this gap in our knowledge, we conducted three studies where we optogenetically activated dopamine neurons while rats were learning associative relationships, both with and without reward. In each experiment, the antecedent cues failed to acquired value and instead entered into value-independent associative relationships with the other cues or rewards. These results show that dopamine transients, constrained within appropriate learning situations, support valueless associative learning.

## Introduction

Dopamine neurons have been famously shown to fire to unexpected rewards ^1^. The most popular idea in the field is that these transient bursts of activity (and the brief pauses on omission of an expected reward) act as the prediction error described in model-free reinforcement learning algorithms^2^. This error term captures the difference between the actual and expected value of the rewarding event and then assigns this ‘excess’ value to any antecedent cue, context, or event ^3^. Critically, such ‘cached’ or learned values are separate from associative or model-based representations linking the actual external events to one another (e.g. the representation linking the cue to the specific sensory features of the reward)^4^. The proposal linking transient changes in the firing of dopamine neurons to value learning is supported mainly by correlative studies, too numerous to mention fully ^(see 5 for full review)^. However support also comes from a growing number of causal reports, showing that artificially induced dopamine transients, with a brevity and timing similar to the physiological prediction-error correlates, are able to drive enduring changes in behavior to antecedent cues, contexts, or events ^6–12^. These changes are assumed to reflect value learning, however almost none of these studies investigate the informational content of the learning to confirm or refute this assumption.

Two studies stand as exceptions to this general statement ^6, 8^. In both, artificially induced dopamine transients, constrained within blocking paradigms to eliminate naturally occurring teaching signals, were shown to cause associations to form between external events. In one study, the association was formed between a cue and the sensory properties of the reward ^8^, and in the other, the association was formed between a cue and the sensory properties of another cue ^6^. Critically, in each experiment, the learning driven by the artificial transient was similar to the learning that would have been observed in the absence of blocking. However, while these results cannot be explained by cached value learning mechanisms, neither of these studies included an assessment of value that was independent of the associative prediction of reward. Thus, these studies did not directly address whether the dopamine transient acted only to facilitate associative learning or also promoted model-free value caching.

In fact, to the best of our knowledge, there is currently no report that directly tests whether a transient increase in the activity of midbrain dopamine neurons functions to assign value directly to cues *in the context of learning*. Here we conducted three experiments to correct this. In each experiment, we used conditioned reinforcement to assess cue value independent of associative predictions about reward, and each time we found that a dopamine transient supported the development of associative representations without endowing the antecedent cues with any value. These data suggest that when dopamine transients occur in a context appropriate for associative learning, they do not act as model-free prediction errors to support cached value learning.

## Results

### Dopamine transients can unblock sensory-sensory learning without endowing the sensory cues with value

Prior to training, all rats underwent surgery to infuse virus and implant fiber optics targeting the ventral tegmental area (VTA; Figure 1). We infused AAV5-EF1α-DIO-ChR2-eYFP (channelrhodopsin-2 (ChR2) experimental group; *n*=8) or AAV5-EF1α-DIO-eYFP (eYFP control group, *n*=8) into the VTA of rats expressing Cre recombinase under the control of the tyrosine hydroxylase (TH) promoter ^13^. After surgery and recovery, rats were food deprived and then trained on the blocking of sensory preconditioning task (Figure 2) used previously ^6^. The use of a design that isolates dopamine transients when rats are learning about neutral information is advantageous because it dissociates the endogeonous value signal elicited by a motivationally-significant reward (e.g. food) from any value that may be inherent in the dopamine transient itself. Thus, it provides an ideal test for determining whether dopamine tranisents endow cues with value.

**Figure 1.**
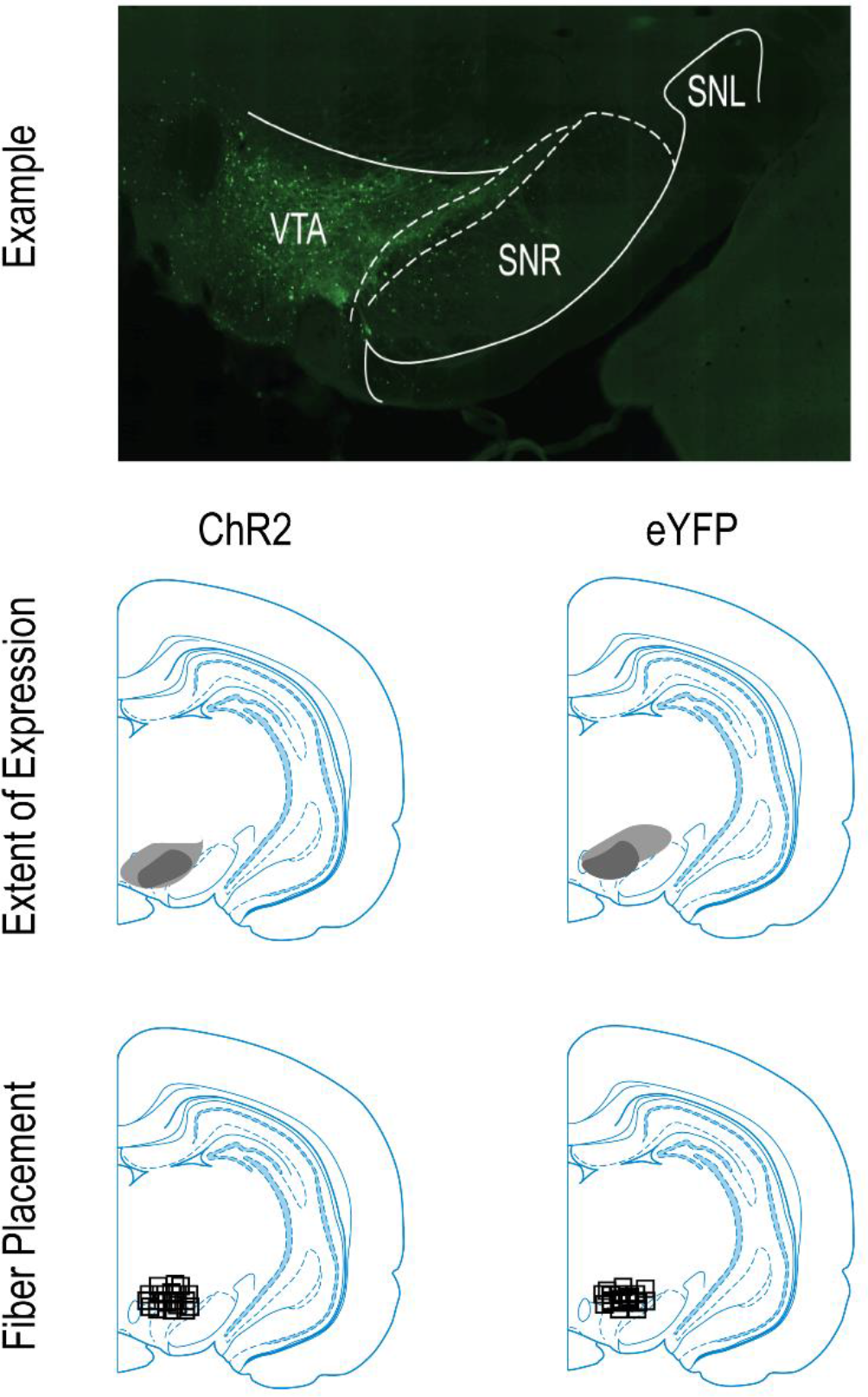
Histological verification of Cre-dependent ChR2 and eYFP in TH+ neurons and fiber placement in the VTA for Experiments 1, 2, and 3. Top row: example of the extent and location of virus expression in VTA, ~ 5 mm posterior to bregma. Middle row: Representation of the bilateral viral expression; dark shading represents the minimal and the light shading the maximal spread of expression at the center of injection sites. Bottom row: Approximate location of fiber tips in VTA, indicated by black squares.

**Figure 2.**
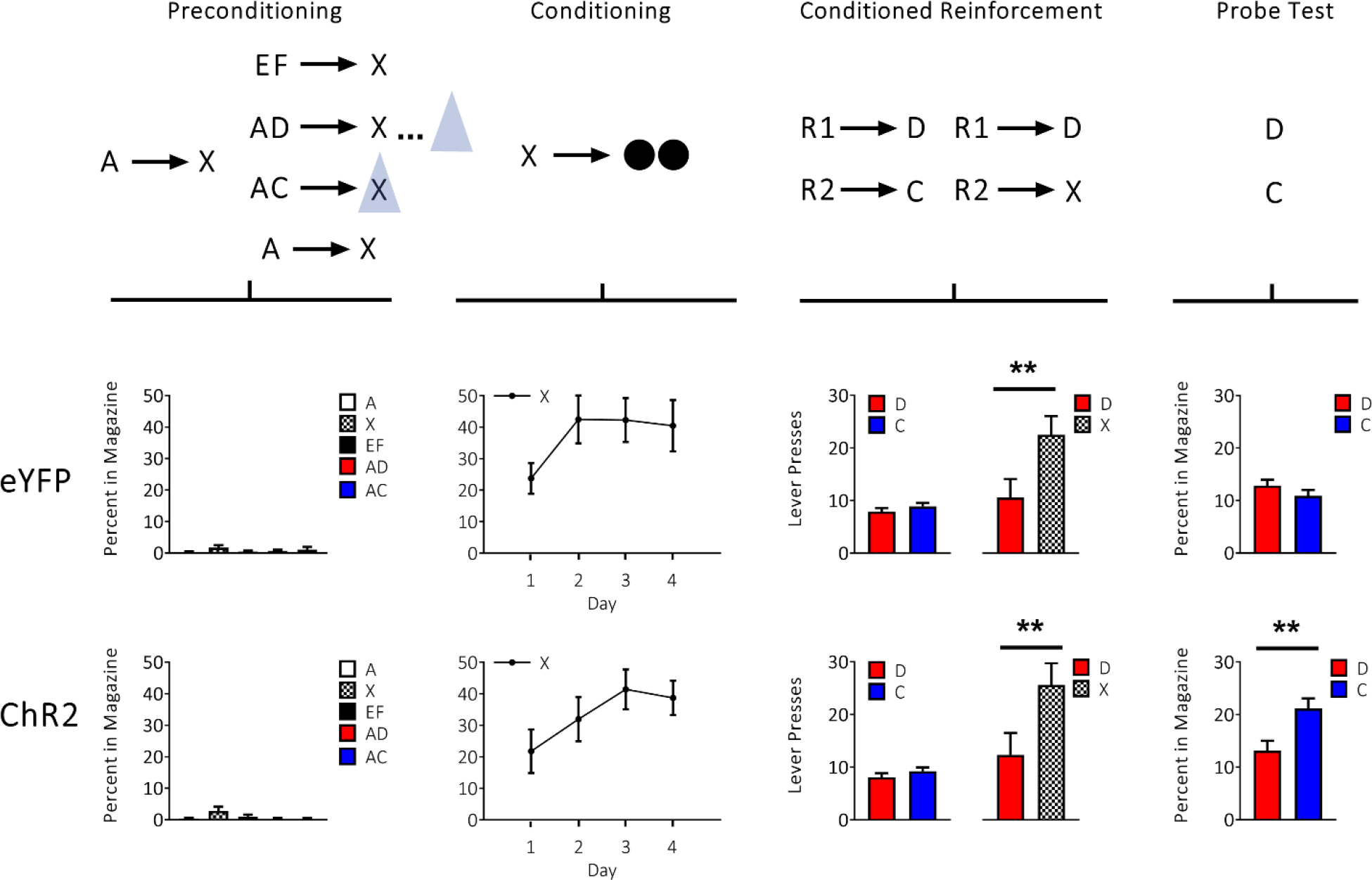
Brief optogenetic stimulation of dopamine neurons in the VTA produces learned associations without endowing cues with cached value. Top row: Schematic illustrating the task design, which consisted of preconditioning and conditioning, followed by conditioned reinforcement and probe testing. A and E indicate visual cues, while C, D, F and X indicate auditory cues. R1 and R2 indicate differently positioned levers, and black circles show delivery of food pellets. VTA dopamine neurons were activated by light delivery in our ChR2 experiment group (*n*=8) but not in our eYFP control group (*n*=8), illustrated by the blue triangle, for 1s at the beginning of X on AC trials and in the inter-trial interval on AD trials. Middle and bottom rows: Plots show rates of responding (±SEM) across each phase of training, aligned to the above schematic. The top row of plots show data for the eYFP control group and bottom row of plots show data for the ChR2 experimental group. For individual rats’ responses see Supplementary Figure 2.

Training began with 2 days of preconditioning. On the first day, the rats received 16 pairings of two 10-s neutral cues (A→X). On the second day, the rats continued to receive pairings of the same two neutral cues (A→X; 8 trials). In addition, on other trials, the first cue, A, was presented together with a second, novel neutral cue, still followed by X (either AC→X or AD→X; 8 trials each). Because A predicts X, this design causes acquisition of the C→X and D→ X associations to be blocked. On AC trials, blue light (473nm, 20Hz, 16-18mW output; Shanghai Laser & Optics Century Co., Ltd) was delivered for 1s at the start of X to activate VTA dopamine neurons, giving the transient an opportunity to both assign value to cue C and also to unblock acquisition of the C→X association. As a temporal control for nonspecific effects, the same light pattern was delivered in the inter-trial interval on AD trials, 120-180s after termination of X. Finally, as a positive control for normal learning, the rats also received pairings of two novel 10-s cues with X (EF→X; 8 trials). As no rewards were delivered, rats in both groups exhibited little responding at the food port on either day of training (Figure 2, preconditioning), with a two-factor ANOVA on the percent of time spent in the food port during cue presentation revealing no significant effects involving group (group: *F*_1,14_=0.027, *p*=0.873; cue × group: *F*_4,56_=0.509, *p*=0.729).

Following preconditioning, the rats underwent 4 days of conditioning to associate X with food reward. To be clear, conditioning is done to provide a behavioral response to assess whether any associations with X were formed during preconditioning. Each day, the rats received 24 trials in which X was presented, followed by delivery of two 45-mg sucrose pellets (X→2US). Rats in both groups acquired a conditioned response, increasing the amount of time spent in the food port during presentation of cue X with no difference in either the level of this response or in the rate at which it was acquired in the two groups (Figure 2, conditioning). A two-factor ANOVA (day × group) revealed only a main effect of day (day: *F*_3,42_=8.258, *p*=0.000; group: *F*_1,14_=0.256, *p*=0.621; day x group: *F*_3,42_=0.513, *p*=0.675).

After conditioning, we assessed whether the preconditioned cue C had acquired any value by virtue of its pairing with a dopamine transient. To do this, we measured the ability of C to promote conditioned reinforcement, which is an iconic test typically used to assess the value of a cue after pairing with a natural reward or drug of abuse ^14^. Critically, conditioned reinforcement is generally accepted as reflecting the value of the cue, independent of any expectations of reward delivery, since it is not normally affected by devaluation of the predicted reward ^15^. In our conditioned reinforcement test, we measured lever pressing during a 30 minute session in which two levers were made available in the experimental chamber, one that led to a 2s presentation of D and another that led to a 2s presentation of C (R1→D, R2→C). In addition, as a positive control for conditioned reinforcement, the same rats also received a second conditioned reinforcement test in which two levers were again made available in the experimental chamber, this time leading to presentation of either D or X (R1→D, R2→X). Because X had been directly paired with reward, it should have acquired value that would support conditioned reinforcement ^15–17^.

The results of the conditioned reinforcement test showed that the dopamine transient did not endow the preconditioned cue C with value; rats in the ChR2 group pressed at the same low rates for both C and D as did eYFP controls (Figure 2, conditioned reinforcement). A two-factor ANOVA (cue × group) revealed no significant effects (cue: *F*_1,14_=1.094, *p*=0.313; group: *F*_1,14_=0.006, *p*=0.940; cue × group: *F*_1,14_=0.004, *p*=0.952). Importantly, the failure to observe conditioned reinforcement for C was not because of any deficiency in conditioned reinforcement, either in our experimental group or using our experimental design, since the ChR2 rats showed conditioned reinforcement equivalent to that seen eYFP rats when they were given a chance to lever press for X, which had acquired value through direct pairing with reward (Figure 2, conditioned reinforcement). A two-factor ANOVA (cue × group) revealed a significant main effect of cue (*F*_1,14_=5.297, *p*=0.037) without any effects of group (group: *F*_1,14_=0.120, *p*=0.734; interaction: *F*_1,14_=0.014, *p*=0.907). Parenthetically, the absence of any group effects suggests that the dopamine transient also failed to add to the value that X acquired by its pairing with reward; this is also suggested by the similar learning rates supported by X during conditioning.

Finally, to confirm that the dopamine tranisent was effective in producing learning between C→X, we assessed the Pavlovian response elicited by the preconditioned cues. To do this, rats received a probe test in which preconditioned cues D and C were presented in an interleaved and counterbalanced manner, alone and without reward, and we measured how much time the rats spent in the food port. Unlike conditioned reinforcement, food port responding in this context is a measure shown previously to reflect a specific expectation of reward delivery, since it is sensitive to devaluation of the predicted reward ^6^. The results of the probe test confirmed that the dopamine transient unblocked learning in the preconditioning phase; rats in the ChR2 group showed a significant increase in the time spent in the food port during presentation of C relative to D, a difference that was not seen in the eYFP group (Figure 2, probe test). A two-factor ANOVA (cue × group) revealed a significant interaction (*F*_1,14_=5.060, *p*=0.041), due to a significant difference between cues C and D in the ChR2 group (*F*_1,14_=6.578, *p*=0.022) but not in the eYFP control group (*F*_1,14_=0.380, *p*=0.547). Thus the introduction of a dopamine transient at the beginning of X on AC trials was sufficient to selectively unblock learning about the C-X association (refer to Supplementary Figure 1 for analyses of this effect with number of entries into magazine) ^6^, but did not endow cue C with model-free value.

### Dopamine transients can accelerate overlearning of sensory-sensory associations without endowing the sensory cues with value

The above experiment demonstrates that a dopamine transient does not endow cues with value, as indexed by an inability of this cue to support conditioned reinforcement. This was despite an effective role in driving the formation of an associative model of events, revealed by the increase in Pavlovian responding at the food cup. This result is contrary to predictions of the hypothesis that the dopamine transient functions as the prediction error described in model-free reinforcement learning algorithms. However this design could be viewed as a weak test of this hypothesis, since value and learning are acting in the same direction. Perhaps C did acquire value, but the conditioned reinforcement test was not sensitive enough to isolate this value. Additionally, it is possible that the desire to respond for food somehow suppresses or interferes with the conditioned reinforcement assessment. To address these possibilities, we designed a more stringent test in which the outcomes predicted by the cached value hypothesis are in opposition to those predicted by associative learning.

To do to this, we returned to the standard sensory preconditioning task ^18^. Sensory preconditioning task is a particularly interesting task to use for this purpose, since responding in the probe test is known to be sensitive to the number of pairings of the cues in the initial preconditioning phase (A→B), increasing with several pairings but then declining thereafter ^19–22^. We observed this effect ourselves, when we exposed rats to 48 pairings of A→B during preconditioning (Supplementary Figure 3), instead of the usual 12 ^6, 17, 23, 24^. Paradoxically, the diminished responding in the probe test with increasing experience in the preconditioning phase is thought to reflect a stronger association between cues A and B, which causes “B-after-A” to become dissociated from “B-alone”^21^. As a result of this, the association between B and reward, acquired in conditioning, does not generalize to the representation of B evoked by the preconditioned cue, A ^19–22^. This phenomenon leads to a somewhat counterintuitive prediction, which is that if the dopamine transient applied in the first experiment is mimicking the normal teaching signal, then delivering it in the standard preconditioning design should diminish evidence of preconditioning, due to an acceleration of this overlearning effect. Of course, this would be in opposition to the value hypothesis, which would simply predict more responding with more value.

**Figure 3.**
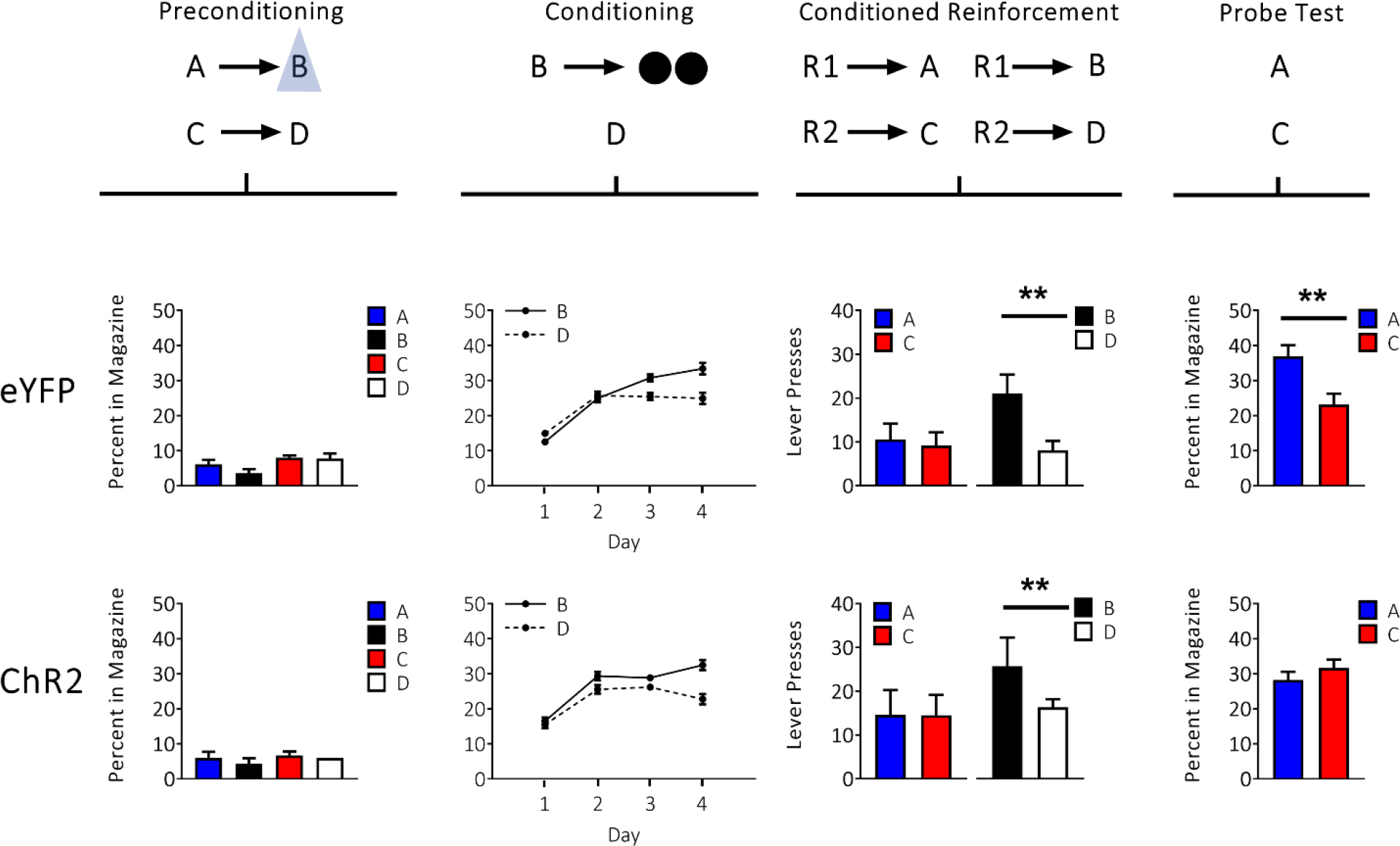
Brief optogenetic stimulation of dopamine neurons in the VTA impairs the sensory preconditioning effect without endowing cues with cached value. Top row: Schematic illustrating the task design, which consisted of preconditioning and conditioning, followed by conditioned reinforcement and probe testing. A, B, C, and D indicate auditory cues, R1 and R2 indicate differently positioned levers, and black circles show delivery of food pellets. VTA dopamine neurons were activated by light delivery, illustrated by the blue triangle, at the beginning of B in our ChR2 group (*n*=14; bottom plots), but in our eYFP control group (*n*=14; top plots). Plots show rates of responding (±SEM) across each phase of training, aligned to the above schematic. For individual rats’ responses, see Supplementary Figure 4.

Prior to training, all rats underwent surgery to infuse virus and implant fiber optics targeting the VTA (Figure 1). We infused AAV5-EF1α-DIO-ChR2-eYFP (ChR2 experimental group; *n*=14) or AAV5-EF1α-DIO-eYFP (eYFP control group, *n*=14) into the VTA of rats expressing Cre recombinase under the control of the TH promoter ^13^. After surgery and recovery, rats were food deprived and then trained on the task. This began with one day of preconditioning, in which the rats received 12 presentations each of two pairs of 10-s neutral cues (i.e. A→B; C→D). On A→B trials, blue light (473nm, 20Hz, 16-18mW output; Shanghai Laser & Optics Century Co., Ltd) was delivered for 1s at the beginning of cue B to activate VTA dopamine neurons. Again, no rewards were delivered during this session, and rats in both groups exhibited little responding at the food port (Figure 3, preconditioning), with a two-factor ANOVA on the percent of time spent in the food port during cue presentation revealing no significant effects involving group (group: *F*_1,26_=0.047, *p*=0.831: cue × group: *F*_3,78_=0.470, *p*=0.704).

Following preconditioning, rats began conditioning, which continued for 4 days. Each day, the rats received 12 trials in which B was presented, followed by delivery of two 45-mg sucrose pellets (B→2US), and 12 trials in which D was presented without reward (D→ nothing). Rats in both groups acquired a conditioned response, increasing the amount of time spent in the food port during cue presentation, and this response came to discriminate between the two cues as conditioning progressed (Figure 3, conditioning). Importantly there was no difference in the level of conditioned responding to B and D across the two groups, or in the rate at which conditioned responding was acquired. Accordingly, a three-factor ANOVA (cue × day × group) revealed main effects of cue (i.e. B vs D; *F*_1,26_=8.692, *p*=0.007) and day (*F*_3,78_=11.979, *p*=0.000) and a significant day × cue interaction (*F*_3,78_=9.783, *p*=0.000), but no significant effects involving group (group: *F*_1,26_=0.018, *p*=0.894; cue × group: *F*_1,26_=0.495, *p*=0.488; day × group: *F*_3,78_=0.271, *p*=0.846; cue × day × group: *F*_3,78_=1.424, *p*=0.242).

Rats then underwent conditioned reinforcement testing (R1→A vs R1→C; R1→B vs R1→D) to assess whether value had accrued to the preconditioned cues. This testing showed that simplifying the preconditioning design and increasing the number of pairings with the dopamine transient was still insufficient to cause acquisition of value by the preconditioned cues. Rats in the ChR2 group lever-pressed at the same low rates for both A and C as controls (Figure 3, conditioned reinforcement), and a two-factor ANOVA (cue × group) revealed no significant effects (cue: A vs. C; *F*_1,26_=0.29, *p*=0.867; group: *F*_1,26_=0.634, *p*=0.433; cue × group: *F*_1,26_=0.019, *p*=0.892). By contrast, rats in both groups showed normal conditioned reinforcement when given a chance to lever press for cue B, which had acquired value through direct pairing with reward (Figure 3, conditioned reinforcement). A two-factor ANOVA (cue × group) revealed a significant main effect of cue (*F*_1,26_=7.132, *p*=0.013) without any group effects (group: *F*_1,26_=0.933, *p*=0.343; cue × group: *F*_1,26_=0.183, *p*=0.672). Again, these data demonstrate that dopamine did not endow the preconditioned cue with value.

Finally, rats received a probe test where we assessed what was learned about the preconditioned cues. In this probe test, we presented cues A and C, alone and without reward, in a counterbalanced and interleaved manner. If the dopamine transient in our design truly functions as a normal teaching signal, then we should see a reduction or loss of food port responding in the probe test, since overlearning of the preconditioned cue pairs normally has this effect (Supplementary Figure 3). Consistent with this prediction, rats in the ChR2 group failed to show the normal difference in food port responding during presentation of A and C in the probe test (Figure 3, probe test). A two-factor ANOVA (cue × group) revealed a significant interaction (*F*_1,26_=4.783, *p*=0.038), due to a difference in responding to A and C in the eYFP group (*F*_1,26_=6.113, *p*=0.020) that was not present in the ChR2 group (*F*_1,26_=0.385, *p*=0.540).

Like the results of the first experiment, these findings show that a dopamine transient is not normally acting as a prediction error that trains cached values; rats would not work to obtain the cue that was paired with the transient. In this experiment, the absence of conditioned reinforcement was observed despite a larger number of presentations of the transient and in the absence of any potential interference due to food predictions that were evident for this cue in the first experiment. Here it is worth emphasizing that while the above results might be hard to square with the cached value hypothesis, they are precisely what would be predicted for associative, model-based learning. That is, sensory preconditioning normally produces a cue that supports model-based food-directed responding but not support conditioned reinforcement, and this responding is sensitive to overtraining. Thus in the above two experiments, the artificial induction of a dopamine transient seemingly supports value-less associative learning rather than cached value learning, when the two are placed in opposition.

### Dopamine transients accelerate configural learning without endowing sensory cues with value

Finally, we tested whether our findings would generalize outside of learning about neutral sensory cues. For this, we used a configural learning task that allowed us to introduce dopamine transients in rats learning about cues and rewards, but in a way that would still allow us to dissociate value-caching from associative learning. On some trials, rats were presented with one of two auditory cues, one predicting reward and one predicting no reward (i.e. A→ nothing; B→2US). On other trials, these same cues were preceeded by a common visual cue, X, and the reward contingencies were switched (i.e. X→A→2US; X→B→ nothing). We delivered a dopamine transient at the onset of A and B, only when these cues were preceeded by X. To learn this task, rats must distinguish “A-alone” and “B-alone” from “X-A” and “X-B” trials. This requires associative representations similar to what is thought to underlie the overlearning effect in sensory preconditioning (Supplementary Figure 3). We reasoned that if dopamine transients were facilitating overlearning in the second experiment by encouraging the formation of such configural representations, then delivering them in the context of this design should facilitate successful discrimination of the compounds. By contrast, if dopamine functions to assign value to antecedent cues, then pairing X with dopamine should simply increase its value, which might interfere with its ability to motivate different responses on X-A versus X-B trials.

The other advantage of this design is that it allows us to present the dopamine tranisent on considerably more trials than was possible during sensory preconditioning. We believe that the sensory preconditioning procedure is appropriate for testing how dopaminergic *prediction errors* normally support learning, since prediction errors that occur in normal learning contexts are precisely timed with regard to external events and disappear relatively rapidly with learning. However, it is plausible that the biological signal – dopamine itself – acts differently when not constrained in this manner. For example, the acquisition of information relevant to the associative model might occur rapidly with relatively little exposure to the transient (as in the first two experiments), whereas acquisition of value may require more exposure (as will be done in this experiment).

Prior to training, all rats underwent surgery to infuse virus and implant fiber optics targeting the VTA (Figure 1). We infused AAV5-EF1α-DIO-ChR2-eYFP (ChR2 experimental group; *n*=6) or AAV5-EF1α-DIO-eYFP (eYFP control group, *n*=6) into the VTA of rats expressing Cre recombinase under the control of the TH promoter ^13^. After surgery and recovery, rats were food deprived and then trained on the task. Configural training continued for 14 days, with rats receiving 40 trials each day. Rats received 10 trials of A without reward, and 10 trials where A was followed by two 45-mg sucrose pellets when immediately preceeded by X (i.e. A→ nothing; X→A→2US). During these sessions, rats also received 10 trials where B was presented with two 45-mg sucrose pellets, and 10 trials in which B was presented without reward when it was immediately preceded by X (i.e. B→2US; X→B→nothing). All four trial types were presented in an interleaved and pseudorandom manner, where no trial type could occur more than twice consecutively. On X→A and X→B trials, blue light (473nm, 20Hz, 16-18mW output; Shanghai Laser & Optics Century Co., Ltd) was delivered for 1s at the beginning of cue A and B to activate VTA dopamine neurons.

Across training, rats gradually learned to discriminate between the differently-rewarded trial types sharing a common cue (Figure 4, configural training). The ability to discriminate these trial types emerged faster in rats in the ChR2 group, demonstrating that dopamine stimulation on the configural trials facilitated learning and did so both when the configural cue predicted reward and also when it did not. A three-factor ANOVA (cue × session × group; Figure 4, top left) revealed a main effect of cue (i.e. AX-A vs. B-BX; *F*_(1,10)_=7.567, *p*=0.020), a main of session (*F*_(6,60)_=7.574, *p*=0.000), and a session × group interaction (*F*_(6,60)_=2.442, *p*=0.035). The source of this interaction was due to rats in the ChR2 group showing faster acquisition of both configural discriminations (i.e. AX-A and X-BX), revealing itself most prominently in the final session block with a significant between-group difference (*F*_(1,10)_=5.166, *p*=0.046). In addition, there was a significant cue × session × group interaction (*F*_(6,60)_=2.470, *p*=0.034), owed to the faster B-XB discrimination relative to XA-A discrimination in ChR2 rats in early trials. However, rats in the ChR2 group showed significant effects of session in regards to both trial types (AX-X: *F*_(6,5)_=8.962, *p*=0.015; B-BX: *F*_(6,5)_=6.205, *p*=0.032), that was not significant in eYFP rats (AX-X: *F*_(6,5)_=0.493, *p*=0.793; B-BX: *F*_(6,5)_=0.335, *p*=0.892). Importantly, these effects were not due to any real-time performance enhancing effect of dopamine, but instead reflected learning. This was evident in a final probe session at the end of configural training in which the laser was not activated on half the trials (Figure 4, probe). During this test, we found no difference in responding to the cues depending on whether or not light was present (light: *F*_(1,10)_=0.077, *p*=0.787; light × group : *F*_(1,10)_=0.039, *p*=0.847; cue × light: *F*_(1,10)_=0.445, *p*=0.520; cue × light × group: *F*_(1,10)_=0.071, *p*=0.795). Thus, activation of dopamine neurons after X enhanced learning to respond differently to the compound cues, suggesting that the dopamine transients facilitated acquisition of the “X-A” and “X-B” configural associations, rather than simply endowing X with a high value.

**Figure 4.**
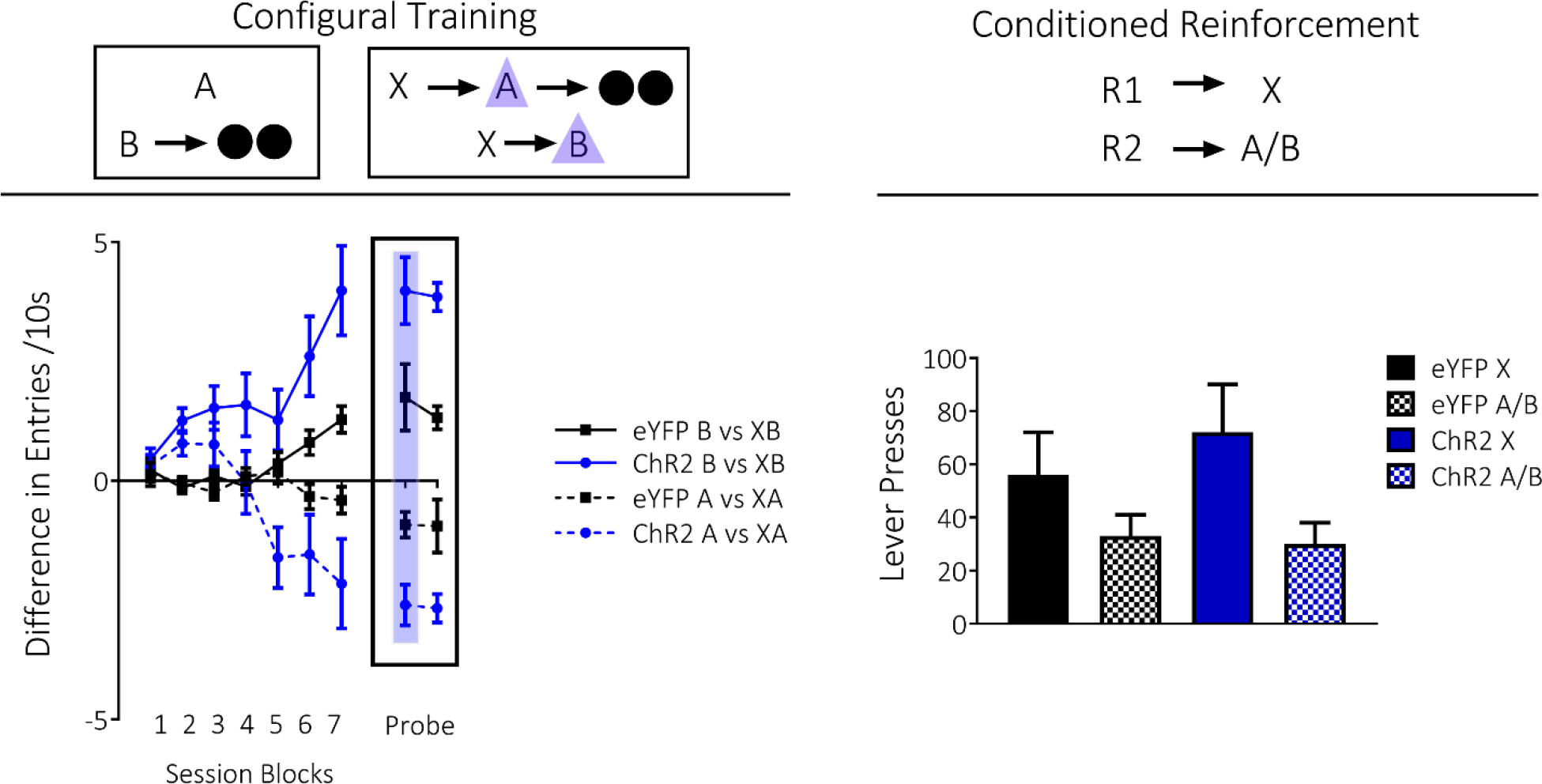
Brief optogenetic stimulation of dopamine neurons facilitates configural learning without endowing cues with value. *Left panel*: Stimulation of dopamine neurons in the ChR2 group (*n*=6) facilitates configural learning, relative to our eYFP control group (*n*=6). A and B indicate auditory cues, X indicates a visual cue, and black circles show delivery of food pellets. VTA dopamine neurons were activated by light delivery (blue triangle) at the beginning of A and B only when preceeded by X. In a final session of configural training (Probe Test, outlined with black rectangle), we presented rats with a standard configural training session, with the exception that half the configural trials were presented with light (shadowed column in blue), and the other half were presented without light. For raw data during configural training, see Supplementary Figure 5. *Right panel*: Stimulation of dopamine neurons during configural training does not produce changes in conditioned reinforcement. R1 and R2 indicate differently positioned levers, and rates of responding are represented as total number of lever presses within a session (±SEM). For individual rats’ responses during conditioned reinforcement testing, see Supplementary Figure 6.

To confirm this, the rats underwent conditioned reinforcement testing to assess whether value had accrued to X, independent of the enhanced discrimination learning. Rats were tested in two sessions in which two levers were made available in the experimental chamber, one that led to a 2s presentation of X and another that led to a 2s presentation of either A or B (R1→X, R2→A/B, with the order of the A and B sessions counterbalanced across subjects in each group). By this time, the rats had received a total of 280 trials where X preceeded dopamine stimulation. If the artificial dopamine transient functioned as a teaching signal for caching value in antecedent cues, as predicted by model-free learning, then X should support robust conditioned reinforcement in the ChR2 group relative to the eYFP group. Yet we found no difference in the rates of lever pressing for X between groups; while both showed higher responding to X relative to A or B, an effect expected based on the strong predictive power of X relative to inconsistent predictors A and B^25, 26^, there was no difference in the magnitude of this difference between groups (Figure 4, conditioned reinforcement). A two-factor ANOVA (cue × group) revealed only a main effect of cue (cue: *F*_(1,10)_=9.608, *p*=0.011; group: *F*_(1,10)_=0.205, *p*=0.660; interaction: cue × group: *F*_(1,10)_=0.829, *p*=0.384). Thus, even when a dopamine transient was delivered many times, we did not see evidence of value caching, but rather we saw evidence for use of the signal to enhance value-independent associative learning.

### Dopamine transients support lever pressing in the absence of associative learning

Lastly, we conducted two additional tests to assess whether the ChR2 rats used in these experiments would press a lever for light delivery into the VTA. This was done to confirm that dopamine neurons were appropriately activated by light in these rats and to show that the rats used here, like those in many other experiments, will engage in behaviors that appear to reflect value when the dopamine neurons are activated in settings that are abnormal or inappropriate for the appearance of a prediction error. In one test, a subset of the ChR2 rats were presented with two levers in the experimental chamber. Pressing one lever resulted in light delivery (473nm, 1s, 50Hz, 16-18mW), while pressing the other had no consequences. Under these conditions, rats exhibited a strong preference for the lever paired with light delivery (*F*_1,6_=27.782, *p*=0.002, Figure 5, left). In a second test, the remaining ChR2 rats and eYFP controls were given the opportunity to press a single lever resulting in light delivery into VTA (473nm, 1s, 50Hz, 16-18mW); two sessions were conducted at 50Hz, and then the frequency was reduced to 20Hz, to test whether the specific frequency used in the experiments was sufficient to maintain responding. Under these conditions, rats in the ChR2 group again pressed at high rates for light delivery into VTA, which was maintained at both frequencies (group: *F*_1,11_= 4.961, *p*=0.048; frequency: *F*_1,11_=3.206, *p*=0.101, *p*=0.101: interaction: *F*_1,11_=1.796, *p*=0.207; Figure 5, right).

**Figure 5.**
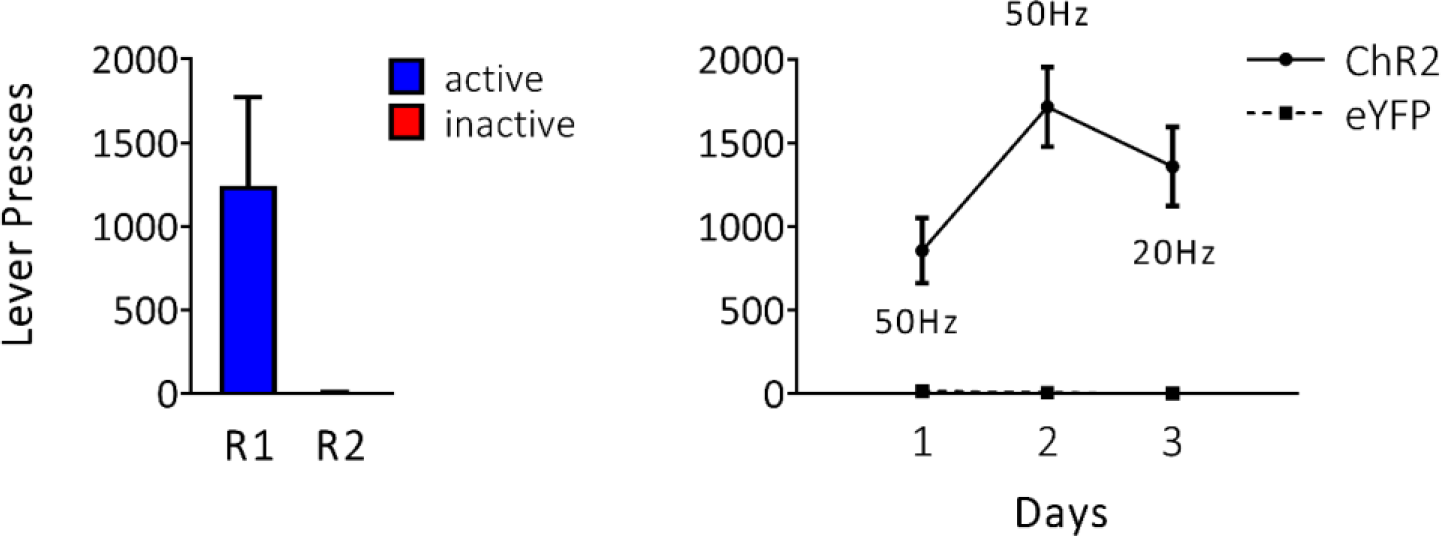
Rats will press a lever to receive direct stimulation of dopamine neurons in the VTA. Rates of responding are represented as the mean level of lever presses (±SEM). Left: Rats from the ChR2 group were given the opportunity to press two differently positioned levers, R1 and R2, where R1 would deliver 50Hz frequency of light delivery into the VTA and R2 did not produce light. Right: A different subset of rats in the ChR2 and eYFP group were given an opportunity to press one lever for 50Hz or 20Hz across different sessions.

## Discussion

Here we have tested whether dopamine transients, like those observed in response to reward prediction errors, obey the predictions of model-free reinforcement learning accounts when placed into contexts appropriate for learning. These theoretical accounts hold that dopaminergic prediction-error signals should result in value caching and should not facilitate the formation of an associative model of events. Recent experiments have challenged these predictions, showing that artificial dopamine transients can support the formation of what appear to be associative representations ^6, 8^. However, these experiments did not directly assess value, independent of their measures of associative learning. This is important because dopamine transients could have two functions, one to support value learning and another to support learning of the associative relationships between external events. Or, as some critics have suggested, the apparent associative learning function in these studies may simply be a side effect of hidden or unappreciated value learning. If either of these explanations are correct, then measures of associative learning and value should covary. Contrary to this prediction, the current results show that dopamine signals, constrained within situations appropriate for learning, cause the acquisition of an associative model of the environment without any evidence of value caching.

Evidence that dopamine transients support valueless associative learning was shown in three experiments. In each experiment, the cues antecedent to the dopamine transient failed to show any evidence of accrued value. In the first two experiments pairing neutral cues with dopamine stimulation did not allow these cues to produce conditioned reinforcement. Specifically, rats would not press a lever to receive presentation of A or C, showing that they did not become valuable to them. In a third experiment, we found that pairing reward-predictive cues with dopamine stimulation also did not alter their ability to produce conditioned reinforcement. That is, rats in the ChR2 group would not press more for X, relative to the eYFP group, even though this cue had been paired with dopamine in the context of reward learning. Importantly, in this experiment rats received a total of 280 trials with dopamine stimulation. Thus, if dopamine was capable of endowing cues with value, we would expect to see a robust increase in the ability of these cues to promote conditioned reinforcement. In contrast to this hypothesis, we found no between-group difference in the ability of these cues to promote conditioned reinforcement. These data show that transient stimulation of dopamine does not endow cues with model-free value.

At the same time, the dopamine transient appeared to facilitate whatever associative or model-based learning was appropriate for these cues. Notably, this was true even when that learning was in direct opppostion to what would be predicted if the dopamine transient were being used for learning value. In the first experiment, the introduction of a dopamine transient at the beginning of X when it was preceded by C unblocked the association between C and X. This unblocking was evident in the increase in food-port responding elicited by C after its associate, X, was paired directly with reward. This result supports prior reports showing associative learning can occur under the influence of an artificial dopamine transient ^6, 8^. The second experiment extended this finding to show that the dopamine transient produces the same effect caused by overlearning of the cue pairs, previously seen both in other labs ^19–22^ and in our hands (see Supplementary Figure 3). Rather spectacularly, here the dopamine tranisent functioned to *reduce* a Pavlovian response directed towards a dopamine-paired cue, in opposition to a cached-value learning hypothesis. Finally, we also showed that introduction of a dopamine transient facilitated configural learning. Again, this was in the absence of any evidence for value caching, which would only have interfered with the configural learning. Thus, these results add to the evidence that dopamine transients are sufficient to instantiate learning of associative representations of events and further show that when placed in contexts where such learning is appropriate, the same transients do not necessarily function to attach a scalar value to those same events as required by current proposals that suggest that dopamine transients function as cached-value teaching signals in model-free reinforcement learning models.

Interestingly, the same rats would work to directly activate the dopamine neurons. This is as expected and suggests that our negative results are not due to poor viral expression, fiber placement or other factors that might impair activation of the dopamine neurons. In addition, this result allows us to emphasize that our findings do not contradict the many other roles that dopamine has been shown to play when delivered for longer time periods or in situations less structured or constrained to isolate associative learning. Indeed, direct activation of dopamine neurons outside of a learning context clearly produces a motivation and behavioral drive that could be called value ^7, 27–32^. In fact, it has recently been shown that directly pairing a cue with optogenetic activation of dopamine neurons, many times and without an obvious external precipitating event, can produce a cue that supports conditioned reinforcement ^27^. These results and our findings here are not mutually exclusive, rather they are complementary, revealing what the biological signal might do under different external conditions.

This point is best illustrated by the consequences it may have in the context of psychological disorders. For example, people with schizophrenia show spurious or overly prominent prediction-error signals, presumed to be dopaminergic, across the course of learning ^33, 34^. Such signals, often still triggered by external events, might drive abnormal or inappropriate associative learning, contributing to the positive symptoms that characterize the disorder, with minimal impact on value learning. On the other hand, addictive drugs typically cause dopamine to be released at times less precisely related to external events and for durations much longer than is typically observed in response to prediction errors. Such release might endow cues, contexts, or events that happen to be present with excessive value, contributing to the aberrant drug-seeking behaviors that characterize the disorder, with minimal impact on associative representations ^35^. These results stress that an understanding of the role of dopamine must consider the form of the signal and the context in which it occurs.

## Online methods

### Surgical procedures

Rats received bilateral infusions of 1.2 μL AAV5-EF1α–DIO-ChR2-eYFP (*n*= 28) or AAV5-EF1α-DIO-eYFP (*n* = 28) into the VTA at the following coordinates relative to bregma: AP: −5.3 mm; ML: ± 0.7 mm; DV: −6.5 mm and −7.7 (females) or −7.0 mm and −8.2 mm (males). Virus was obtained from the Vector Core at University of North Carolina at Chapel Hill (UNC Vector Core). During surgery, optic fibers were implanted bilaterally (200-μm diameter, Thorlabs) at the following coordinates relative to bregma: AP: −5.3 mm; ML: ± 2.61 mm and DV: −7.05 mm (female) or −7.55 mm (male) at an angle of 15° pointed toward the midline.

### Apparatus

Training was conducted in eight standard behavioral chambers (Coulbourn Instruments; Allentown, PA), which were individually housed in light- and sound-attenuating boxes (Jim Garmon, JHU Psychology Machine Shop). Each chamber was equipped with a pellet dispenser that delivered 45-mg pellets into a recessed food port when activated. Access to the food port was detected by means of infrared detectors mounted across the opening of the recess. Two differently shaped panel lights were located on the right wall of the chamber above the food port. The chambers contained a speaker connected to white noise and tone generators and a relay that delivered a 5-kHz click stimulus. A computer equipped with GS3 software (Coulbourn Instruments, Allentown, PA) controlled the equipment and recorded the responses. Raw data were output to and processed in Matlab (Mathworks, Natick, MA) to extract relevant response measures, which were analyzed in SPSS software (IBM analytics, Sydney, Australia).

### Housing

Rats were housed singly and maintained on a 12-h light–dark cycle; all behavioral experiments took place during the light cycle. Rats had *ad libitum* access to food and water unless undergoing the behavioral experiment, during which they received either 8 grams or 12 grams of grain pellets- for females and males, respectively-daily in their home cage following training sessions. Rats were monitored to ensure they did not drop below 85% of their initial body weight across the course of the experiment. All experimental procedures were conducted in accordance with the NIDA-IRP Institutional Animal Care and Use Committee of the US National Institute of Health guidelines.

### General behavioral procedures

Trials consisted of 10-s cues as described below. Trial types were interleaved in miniblocks, with the specific order unique to each rat but counterbalanced within each group. Intertrial intervals varied around a 6-min mean. Unless otherwise noted, daily training was divided into a morning (AM) and afternoon (PM) session.

### Response measures

We measured entry into the food port to assess conditioned responding. Food port entries were registered when the rat broke a light beam placed across the opening of the food port. This simple measure allowed us to calculate a variety of metrics including response latency after cue onset, number of entries to the food port during the cue, and the overall percentage of time spent in the food port during the cue. These metrics were generally correlated during conditioning, and all reflect to some extent the expectation of food delivery at the end of the cue in a task such as that used here. Generally when analyzing behavior during the sensory preconditioning task, we measure conditioned responding by calculating the percent of time rats spend in the food port during cue presentation^17, 23, 24^. The exception to this has been when analyzing behavior in the blocking of sensory preconditioning task. Specifically, in our previous manuscript we reported behavior in the blocking of sensory preconditioning procedure using number of entries into the food port during cue presentation^6^. However, in the current manuscript we have represented the data as the percent of time spent in the food port to be consistent across experiments within this manuscript. Importantly, both measures of behavior were highly correlated and elicited significant results in our critical probe tests (see Supplementary Figure 1 for data as assessed by entries into the food port in Experiment 1, as well as additional analyses comparing the two response measures for this experiment).

### Histology

All rats were euthanized with an overdose of carbon dioxide and perfused with phosphate buffered saline (PBS) followed by 4% paraformaldehyde (Santa Cruz Biotechnology Inc., CA). Fixed brains were cut in 40-μm sections to examine fiber tip position and virus expression under a fluorescence microscope (Olympus Microscopy, Japan).

### Statistical analyses

All statistics were conducted using the SPSS 24 IBM statistics package. Generally, analyses were conducted using a mixed-design repeated-measures ANOVA. All analyses of simple main effects were planned and orthogonal and therefore did not necessitate controlling for multiple comparisons. Data distribution was assumed to be normal, but homoscedasticity was not formally tested. Except for histological analysis, data collection and analyses were not performed blind to the conditions of the experiments.

### Data availability

The data that support the findings of this study, and any associated custom programs used for its acquisition, are available from the corresponding authors upon reasonable request.

## Experiment 1

### Subjects

Sixteen experimentally naive male and female Long-Evans transgenic rats of approximately 4 months of age at surgery and carrying a TH-dependent Cre expressing system (NIDA Animals Breeding Facility) were used in this study. Sample sizes were chosen based on similar prior experiments that have elicited significant results with a similar number of rats. No formal power analyses were conducted. Rats were randomly assigned to groups and distributed equally by age, gender and weight. Prior to final data analysis, four rats were removed from the experiment due to virus or cannula misplacement as verified by histological analysis.

### Blocking of sensory preconditioning

This procedure was essentially identical to that used previously ^6^. Briefly, training used a total of six different stimuli, drawn from stock equipment available from Coulbourn and included four auditory (tone, siren, clicker, white noise) and two visual stimuli (flashing light, steady light). Assignment of these stimuli to the cues depicted in Figure 2 and described in the text was counterbalanced across rats in each group within each modality (A and E were visual while C, D, F and X were auditory).

Training began with 2 d of preconditioning. On the first day, the rats received 16 presentations of A→X, in which a 10-s presentation of A was immediately followed by a 10-s presentation of X. On the second day, the rats received 8 presentations of A→X alone, as well as 8 presentations each of three 10-s compound cues (EF, AD, AC) followed by X (i.e., EF→X; AD→X; AC→X). On AC trials, light (473 nm, 16–18 mW output; Shanghai Laser & Optics Century Co., Ltd) was delivered into the VTA for 1 s at a rate of 20 Hz at the beginning of X; on AD trials, the same light pattern was delivered during the intertrial interval, 120–180 s after termination of X. Following preconditioning, rats underwent 4 d of conditioning in which X was presented 24 times each day and was followed immediately by delivery of two 45-mg sucrose pellets (5TUT; TestDiet, MO).

Following this training, rats received two different test sessions: a conditioned reinforcement test to assess value attribution to the preconditioned cues and a probe test to provide formal evidence of preconditioning. The order of these tests sessions was counterbalanced within the rats in each group such that half the rats in each group received the conditioned reinforcement test first, and the other half of the rats received the probe test first. On the following day, this order was reversed so that all rats received the alternate test. This pattern was then repeated once more so that all rats had two conditioned reinforcement tests and two probe tests. During conditioned reinforcement testing, the food ports were removed from the chamber and two levers were placed into the box for the first time, as in previous experiments ^16, 17^. Pressing one lever resulted in a 2s presentation of cue D, while pressing the other lever resulted in a 2s presentation of cue C. This session lasted for 30 minutes. During the probe test, the chambers were kept as per the earlier training sessions and cues C and D were presented six times each in an interleaved and counterbalanced order, alone and without reward. Analyses were conducted on data over all sessions, where analyses for the probe tests were restricted to the last 5s of the first six trials of cue presentation in each probe session as done previously^6^.

Following these test sessions, all rats received a final conditioned reinforcement test session. During this session, the two levers were put into the chamber and pressing one again resulted in a 2s presentation of cue D, while pressing the other resulted in a 2s presentation of cue X. These sessions lasted for 30 minutes. This provided a positive control to show that these rats would exhibit conditioned reinforcement for a cue directly paired with food using our procedures.

## Experiment 2

### Subjects

Twenty-eight experimentally-naive male and female Long-Evans transgenic rats of approximately 4 months of age at surgery and carrying a TH-dependent Cre expressing system (NIDA animal breeding facility) were used in this study. Sample sizes were chosen based on similar prior experiments that elicited significant results with a similar number of rats. No formal power analyses were conducted. Rats were randomly assigned to groups and distributed equally by age, gender and weight. Prior to final data analysis, five rats were removed from the experiment due to illness, virus or cannula misplacement.

### Sensory preconditioning

Training used a total of four different auditory stimuli, drawn from stock equipment available from Coulbourn, which included tone, siren, clicker and white noise. Assignment of these stimuli to the cues depicted in Figure 3 and described in the text was counterbalanced across rats. Training began with 1 d of preconditioning, in which where rats received 12 presentations of the A→B serial compound and 12 trials of the C→D serial compound. Following preconditioning, rats began conditioning, in which they received 24 trials of B and 24 trials of D, where B was immediately followed by presentation of two 45-mg sucrose pellets and D was presented in the absence of reward. Rats received 4 days of conditioning in this manner.

Following this training, rats again received a probe test to assess preconditioning and a conditioned reinforcement test to assess value attribution to the preconditioned cues. The order of these test sessions was again counterbalanced so that half the rats in each group received the conditioned reinforcement test first and the other half of the rats received the probe test first. During the probe test, the chambers were kept as per the earlier training sessions and cues A and C were presented six times each in an interleaved and counterbalanced order, alone and without reward. During the conditioned reinforcement test, the food ports were removed from the chamber and two levers were placed in the box for the first time, as done in previous experiments ^16, 17^. Pressing one lever resulted in a 2s presentation of cue A, while pressing the other lever resulted in a 2s presentation of cue C. These sessions lasted for 30 minutes.

Following these test sessions, all rats received a final conditioned reinforcement test session. During this session, the two levers were put into the chamber and pressing one again resulted in a 2s presentation of cue B, while pressing the other resulted in a 2s presentation of cue D. These sessions lasted for 30 minutes and provided a positive control for the ability of thease rats to show conditioned reinforcement under our procedures.

## Experiment 3

### Subjects

Twelve experimentally-naive male and female Long-Evans transgenic rats of approximately 4 months of age at surgery and carrying a TH-dependent Cre expressing system (NIDA animal breeding facility) were used in this study. Sample sizes were chosen based on similar prior experiments that elicited significant results with a similar number of rats. No formal power analyses were conducted. Rats were randomly assigned to groups and distributed equally by age, gender and weight. All rats were included in the final analyses.

### Configural Training

Training used a total of three different stimuli, a flashing light controlled by Coulbourn, and a chime and siren sound produced by Arduino software. Assignment of these stimuli to the cues depicted in Figure 4 and described in the text was counterbalanced across rats (X was visual, while A and B were auditory). Configural training consisted of four different trial types within the same session, randomly presented, where no trial type could occur more than twice consecutively. On elemental trial types, A was presented without food, while B was presented and followed immediately by delivery of two 45 -mg sucrose pellets. On configural trials, X was presented immediately before A and B to form two serial compounds X→A and X→B. In contrast to elemental trials, X→A was followed immediately by two 45-mg sucrose pellets, while X→B was presented in the absence of reward. Rats received 14 days of training, receiving one session consisting of 40 trials per day. In a final session, we gave rats a standard training session with the exception that light was omitted on half the trails. All other aspects of the session remained the same as during training.

Following this training, rats received two conditioned reinforcement tests. During these tests, the food ports were removed from the chamber and two levers were placed in the box for the first time, as done in previous experiments. In one conditioned reinforcement test, pressing one lever resulted in a 2s presentation of X, while pressing the other resulted in a 2s presentation of A. In the other conditioned reinforcement test, pressing one lever continued to result in a 2s presentation of X, while pressing the other resulted in a 2s presentation of B. The order in which rats received these sessions was fully counterbalanced. These sessions lasted for 30 minutes.

### Brain Stimulation

Rats received one of two brain stimulation procedures. A subset of rats from the ChR2 group received a choice test in which two levers were again inserted into the chambers and the food ports removed. Pressing one lever resulted in delivery of 1s of light into the VTA (50Hz, 473nm), while pressing the other lever had no effect. Another subset of rats from both the ChR2 and eYFP groups received three sessions where one lever was made available in the chambers. Here, pressing the lever resulted in 1s of light delivery into VTA at a frequency of 50Hz for the first 2 sessions and 20Hz for the third session. Sessions lasted for 30 minutes. As the standard deviation was found to increase with the mean, the data were analyzed using the logarithmic transform log (a+1) ^36, 37^.

