## Supplementary Information for "Dopamine transients delivered in learning contexts do not act as model-free prediction errors"

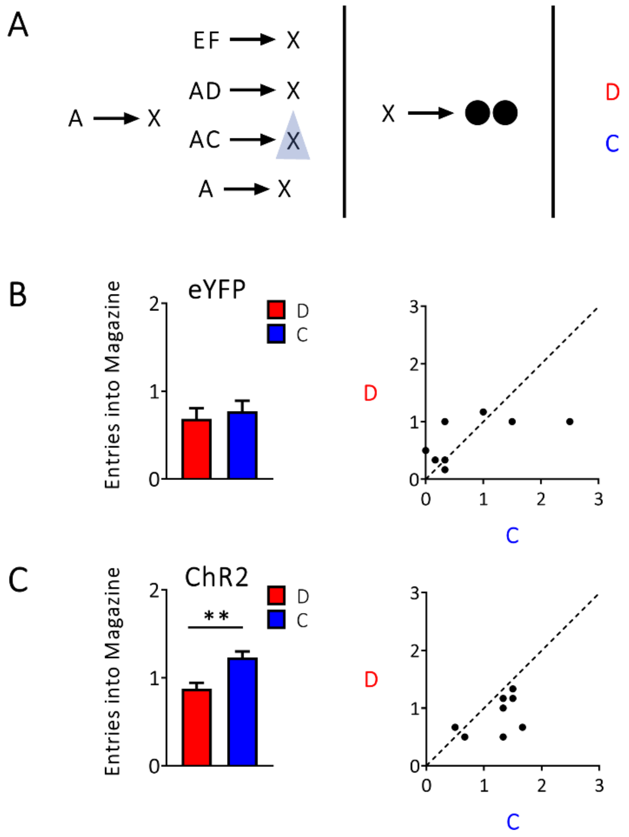


**Supplementary Figure 1. Dopamine transients drive the formation of associations between events without endowing cues with cached value when analyzing number of entries made into the food port.** Rates of responding are represented as the number of entries made into the food port during stimulus presentation (±SEM). A) Experimental Design: Rats were first given preconditioning, where dopamine neurons in the ventral tegmental area were stimulated for 1s at the beginning of X when it was preceded by the AC compound. Then, rats received conditioning where cue X was paired with reward. Finally, rats received a critical probe test where cues C and D were presented alone and without reward. B) Responding during the critical probe test for the eYFP control group (*n*=8), left panels shows mean data for the group, right panel shows individual rats' responses. C) Responding during the critical probe test for the ChR2 group, left panel illustrates mean level of responding for the group, right panel indicates individual rats' responses. As with measures of percent spent in the magazine (reported in the main text), we found that rats in the ChR2 group showed a significantly higher number of entries into the magazine when C was presented relative to D ($F_{1,7}$=6.858, $p$=0.034), while rats in the eYFP group showed no difference in responding during C relative to D ($F_{1,7}$=0.121, $p$=0.738). Further, the correlation between the difference between responding to C and D in terms of the number of entries made into the magazine and the percent of time spent in the magazine was ~55%, and the total variability in the difference in number of responses made towards C and D that could be predicted from the difference in percent of responding in the magazine during C and D was ~30% ($F_{1,14}$=5.514, p=0.034). Thus, the two measures were significantly correlated.

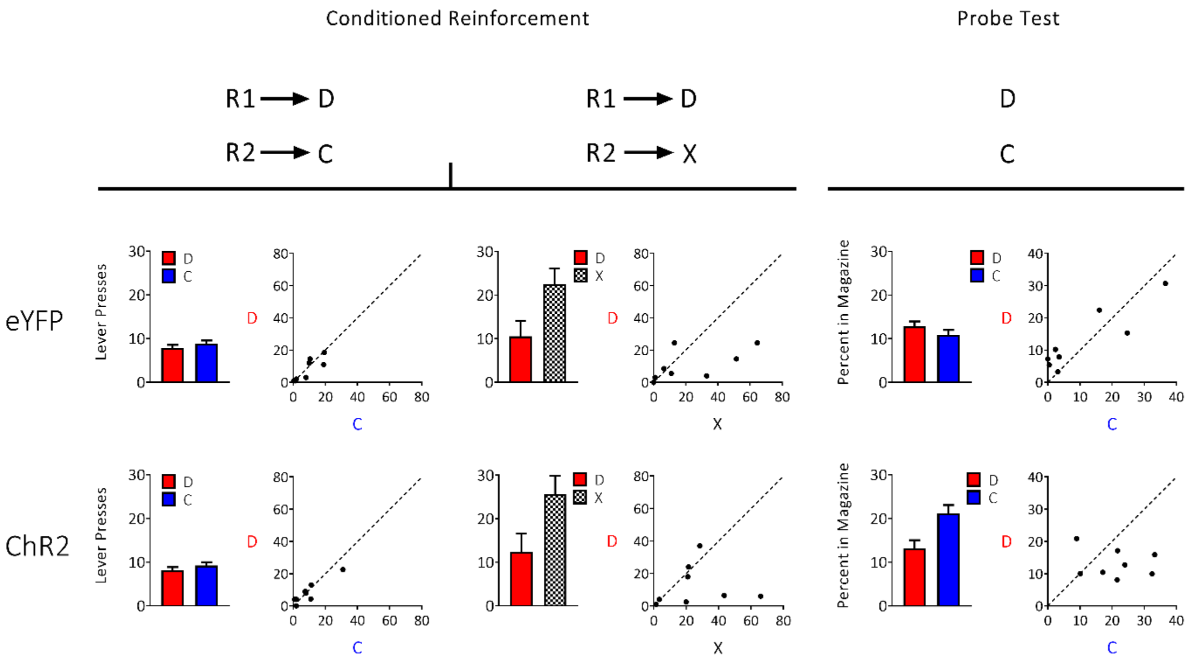


**Supplementary Figure 2. Individual data points from Experiment 1 in main text.** The graphs illustrate the mean level of responding (±SEM; left) and individual rats' responses (right) for the critical tests from Experiment 1: conditioned reinforcement tests with C and D, conditioned reinforcement tests with D and X, and probe tests with C and D. To the point that responding is equal to both cues, the individual points in the scatterplots will congregate around the diagonal.

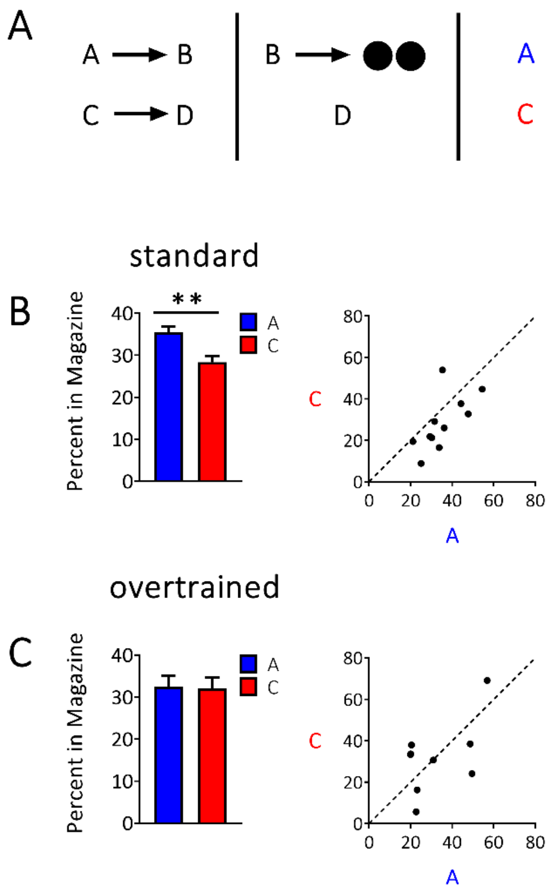


**Supplementary Figure 3. The sensory preconditioning effect is susceptible to overtraining.** Rates of responding are represented as time spent in the food port during stimulus presentation (±SEM). A) Experimental design: rats received either 1 day (group "standard": *n*=11) or 4 days (group "over-trained", *n*=9) of preconditioning prior to commencement of conditioning. Black circles indicate delivery of food pellets. B) Responding to cues A and C in the critical probe test for the standard group where left panel illustrates group mean levels of responding and right panel indicates scatterplots of individual rats' responses. C) Responding in the critical probe test for the over-trained group where again left panel illustrates group mean levels of responding and the right panel indicates scatterplots of individual rats' responses. Here, we found that rats in the standard group showed a significant preconditioning effect. That is, rats in the standard group show a significantly higher percentage of time spent in the food port during presentation of stimulus A relative to C ($F_{1,10}$=5.584, $p$0.040). This was not seen in rats in the over-trained group ($F_{1,8}$=0.007, $p$=0.936). These data showed that the sensory preconditioning effect is susceptible to overtraining, consistent with past research[1-4].

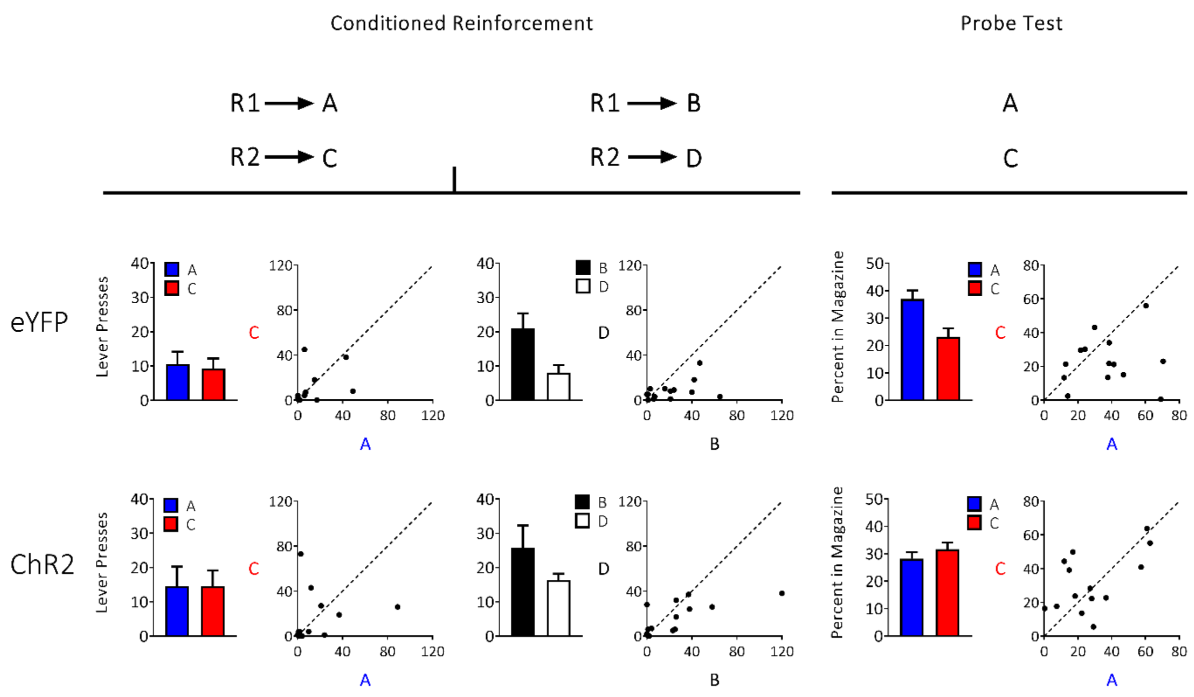


**Supplementary Figure 4. Individual data points from Experiment 2 in main text.** The graphs illustrate the mean level of responding (±SEM; left) and individual rats' responses (right) for the critical tests from Experiment 2: conditioned reinforcement tests with A and C, conditioned reinforcement tests with B and D, and probe tests with A and C. To the point that responding is equal to both cues, the individual points in the scatterplots will congregate around the diagonal.

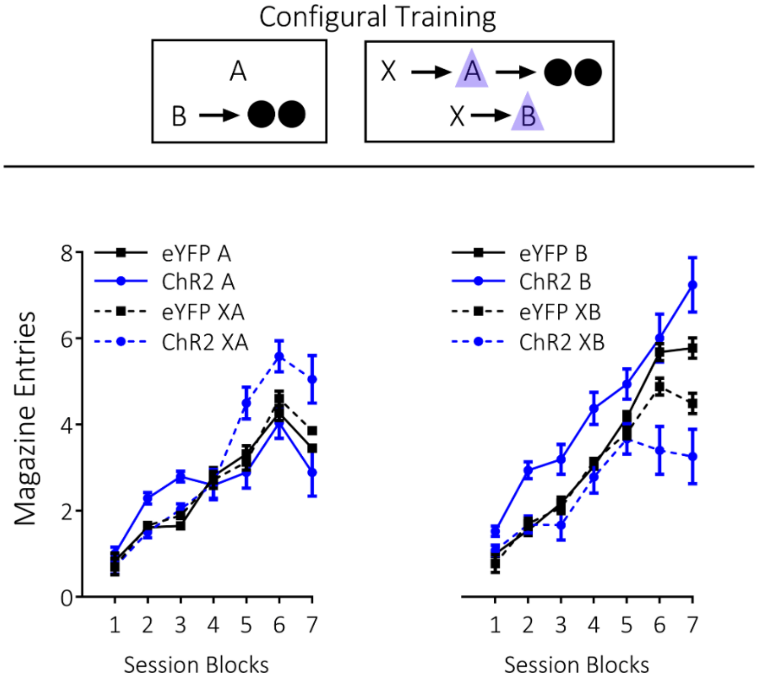


**Supplementary Figure 5. Raw data from Configural Training in Experiment 3.** A and B indicate auditory cues, X indicates a visual cue, and black circles show delivery of food pellets. VTA dopamine neurons were activated by light delivery (blue triangle) at the beginning of A and B only when preceeded by X. The graphs illustrate the mean levels of magazine entries (±SEM) during A and B. Across training rats gradually learned to discriminate between the trial types (Figure 4, top panel), where discrimination between the B trial types (B→2US; X→B→nothing) emerged more quickly than that shown on the A trial types (A→nothing; X→A→2US). Rats in the ChR2 group showed faster discrimination between both trial types, demonstrating that dopamine stimulation on the configural trials facilitated the discrimination between these trial types. This was supported by statistical analyses on the absolute values of the data. A three-factor ANOVA (cue × session × group) on the data from the B trial types (i.e. B vs. XB; Figure 4 top right) revealed a main effect of cue ($F_{(1,10)}$=12.059, $p$=0.006) and a cue × group interaction ($F_{(1,10)}$=5.425, $p$=0.042). This interaction was due to a significant difference in responding between the cues in the ChR2 group ($F_{(1,10)}$=16.830, p=0.002), that was not present in the eYFP group ($F_{(1,10)}$=0.654, $p$=0.438). These analyses also revealed a main effect of session ($F_{(6,60)}$=21.248, $p$=0.000), and a cue by session interaction ($F_{(1,10)}$=5.912, $p$=0.000), driven by a significant main effect of session with regards to B ($F_{(6,5)}$=5.672, $p$=0.038), that did not reach significance in relation to XB ($F_{(6,5)}$=4.238, $p$=0.067). There was no between-group difference in overall levels of responding to the cues ($F_{(1,10)}$=0.121, $p$=0.735). A three-factor ANOVA (cue x session x group) on the data from the A trial types (i.e. A vs. XA; Figure 4 top middle) showed a significant main effect of session ($F_{(6,60)}$=17.577, $p$=0.000), a cue × session interaction ($F_{(6,60)}$=4.888, $p$=0.000), and a cue × session × group interaction ($F_{(6,60)}$=3.769, $p$=0.003). The source of this interaction was due to a significant difference in the responding to the cues in ChR2 group that emerged most prominently in the later sessions ($F_{(1,10)}$=7.383, $p$=0.022), which was not present in the eYFP rats ($F_{(1,10)}$=0.264, $p$=0.618). Again, there were no differences in between-group levels of responding over these sessions ($F_{(1,10)}$=0.289, $p$=0.603).

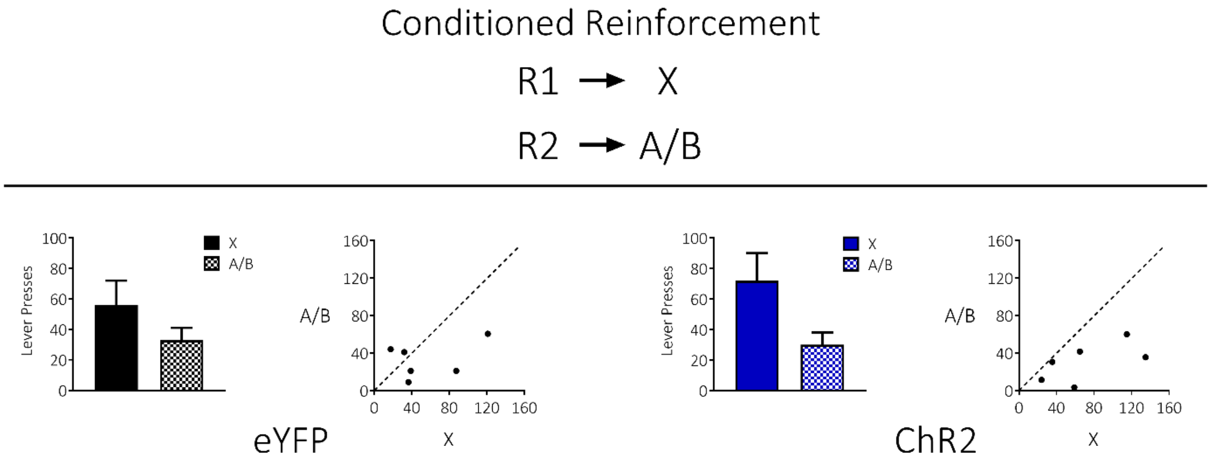


**Supplementary Figure 6. Individual data points from Experiment 3 in main text.** The graphs illustrate the mean level of responding (±SEM; left) and individual rats' responses (right) for the conditioned reinforcement tests. The conditioned reinforcement tests compare responding for X relative to A/B. To the point that responding for both cues is equal, the individual points in the scatterplots will congregate around the diagonal.
